# Associations Between QuantiFERON-TB Gold Plus IFN_γ_ Concentrations and Progression to Symptomatic Tuberculosis in Global High-Burden TB Settings

**DOI:** 10.64898/2026.01.29.702660

**Authors:** Justine Sunshine, Michael Shaffer, Linda L. Han, Deepali Gaikwad, Amelia A. Houana, Maria Tarcela Gler, Sri Rezeki Hadinegoro, Willem A. Hanekom, Javier R. Lama, Monde Muyoyeta, Sissy Musala, Videlis Nduba, Valeria C. Rolla, Tapash Roy, Jayne S. Sutherland, Celso Khosa, Anne Wajja, Timothy M. Walker, Amy Cinar, Alexander C. Schmidt, Alemnew F. Dagnew, Nicole Frahm, TBV02-E01 Study Group

## Abstract

**Introduction:** Predictive biomarkers for symptomatic tuberculosis (TB) progression would transform targeted prevention efforts. Although interferon-gamma release assays (IGRAs), including QuantiFERON® TB-Gold Plus (QFT-Plus), have been studied for this purpose, systematic evaluation of the QFT-Plus TB1 and TB2 Interferon-Gamma (IFNγ) concentrations remains limited, particularly in high-burden TB settings.

**Methods:** Baseline TB1 and TB2 IFNγ concentrations from 5,259 participants in TB-endemic regions were analyzed in relation to subsequent TB outcomes over a median of 525 days follow-up. Participants were categorized as controls (no TB), suspected TB (no microbiological confirmation), or laboratory-confirmed TB, including a subset meeting a stringent case definition (≥2 positive microbiologic tests). Associations between baseline IFNγ concentrations and progression to symptomatic TB were assessed.

**Results:** In the full cohort (IGRA+/-participants), baseline TB2 IFNγ concentrations were significantly higher compared to controls among participants who developed suspected TB (p=0.01), laboratory-confirmed TB (p=0.01), or met the stringent case definition (p<0.0001). In IGRA+ participants, baseline TB2 concentrations were significantly higher than controls in suspected (p=0.01) and laboratory-confirmed (p=0.02) groups. Associations with baseline TB1 IFNγ concentrations and TB progression were observed for participants meeting the stringent case definition within the full cohort (p=0.009). Among stringent definition cases, TB2 concentrations achieved an area under the Receiver Operating Characteristic curve of 0.84, with a sensitivity of 80% and specificity of 78%.

**Conclusions:** Quantitative IFNγ concentrations from QFT-Plus, particularly TB2, were associated with progression to symptomatic TB, met or exceeded WHO-recommended sensitivity and specificity thresholds for predictive biomarkers, and may support biomarker-based stratification in TB clinical research.

**Summary:** In an exploratory analysis from a global tuberculosis (TB) epidemiology study, higher QuantiFERON-TB Gold Plus IFNγ concentrations were associated with progression to symptomatic TB and met WHO predictive biomarker thresholds, supporting their potential value for risk stratification in TB research.

## Introduction

Tuberculosis (TB) remains one of the most significant global health challenges with more than 10 million cases and over one million deaths reported in 2023 [1]. It is estimated that one fourth of the world’s population is or has been infected with *Mycobacterium tuberculosis* (*Mtb*). While many infections are believed to be contained or eliminated, approximately 5–10% of individuals infected with *Mtb* will progress to TB disease, with the highest risk of progression within the first two years following infection [2]. Targeting individuals at the highest risk of developing active TB with preventive treatment is central to the WHO End TB Strategy [3]. A predictive biomarker for progression to active TB would enable more efficient use of resources by identifying individuals most likely to develop disease, supporting both local care and global elimination efforts. Furthermore, as most current TB vaccine and prevention trials use symptomatic, microbiologically confirmed TB as the primary endpoint, there is also interest in identifying biomarkers that predict progression to symptomatic TB disease in a manner that aligns with a trial’s endpoint definition. Such endpoint-matched biomarkers would be valuable in designing efficient, well-powered trials targeting those at highest risk.

The WHO specifies minimum sensitivity and specificity of >75% for tests predicting progression within two years, providing an important benchmark for evaluating biomarkers of symptomatic TB progression[4]. Current diagnostic tests, including the commonly used Interferon-Gamma Release Assay (IGRA), do not currently meet this minimum performance criteria[5, 6]. IGRA tests measure immunoreactivity to TB antigens through detection of released interferon-gamma (IFNγ), generally via an Enzyme-Linked Immunosorbent Assay (ELISA). As such, the IGRA test does not measure *Mtb* burden but rather a TB-specific effector or memory immunological response, which indirectly measures prior or ongoing *Mtb* exposure. A frequently used IGRA test is the QuantiFERON® TB-Gold Plus (QFT-Plus), which contains two TB antigen stimulation tubes: TB1 and TB2 [7]. The TB1 tube contains peptides derived from TB antigens ESAT-6 (Early Secreted Antigen Target 6 kDa) and CFP-10 (Culture Filtrate Protein 10) and is designed to detect TB-specific CD4^+^ T-cell responses [7]. The TB2 tube contains the same peptides as TB1 but with additional shorter peptides designed to detect TB-specific CD8^+^ T-cell responses [7]. Whereas previous iterations of the QFT test (QFT Gold-in-Tube, GIT) only contained the TB1 tube (with additional TB7.7 peptides), the TB2 tube was included in the QFT-Plus test due to emerging literature that showed TB-specific CD8^+^ T-cell responses were associated with increased *Mtb* bacterial load, recent exposure to *Mtb*, and active TB disease[8-10].

While the QFT test qualitative readout (positive/negative/indeterminate) has been shown to be a poor predictive biomarker [11], there has been considerable interest in the evaluation of the quantitative IFNγ readout in predicting progression to TB disease. Most studies of quantitative IFNγ responses used QFT-GIT data and therefore focused on the single antigen tube. A meta-analysis of 37 studies found a concentration-dependent relationship between IFNγ levels and risk of progression to TB disease [12], a finding supported by additional studies in high and low-endemic settings [13, 14]. Notably, across all studies, there was considerable variation in the cohorts as well as substantial overlap in TB-specific IFNγ responses between individuals who do and do not progress to symptomatic TB disease, thus limiting its discriminatory power [12]. In addition, both the QFT-GIT and QFT-Plus tests are prone to considerable analytical variability [15] and thus it is difficult to directly compare results across studies conducted in different laboratories, as changes to sample handling and processing can impact quantitative measurements[16-18]. For the QFT-Plus assay, several studies have affirmed that TB2-specific IFNγ values are higher in individuals with active TB disease [19, 20], but studies have been very limited in evaluating TB1 vs TB2-specific responses in progression to TB disease [13], especially in high-burden countries.

We recently completed a global observational epidemiologic study in 45 sites across 14 countries to evaluate IGRA status among people living in settings of high TB risk [21]. This study was designed to describe both the proportion of IGRA positivity by site and the incidence of suspected and laboratory-confirmed (lab-confirmed) TB to inform and build capacity for phase 3 TB vaccine efficacy trials. IGRA status was assessed using the QFT-Plus test and thus allowed us to systematically evaluate the utility of QFT-Plus quantitative IFNγ concentrations in predicting progression to symptomatic TB disease in multiple real-world, high-burden TB settings under the same clinical and laboratory protocols. Therefore, in this exploratory analysis, we evaluated baseline QFT-Plus quantitative TB1 and TB2 IFNγ concentrations across control, suspected TB, and lab-confirmed groups to assess their association and predictive performance for progression to symptomatic TB disease.

## Methods

### Study Design

An epidemiological study to assess IGRA positivity was conducted from 20 December 2021 through 16 August 2024 (NCT05190146). Detailed study design, procedures, and results are described elsewhere [21]. Briefly, the study enrolled 7,164 participants aged 15–34 years from high-burden TB communities across 14 countries but with no TB diagnosis in the past two years. Study visits occurred every 6 months after screening (baseline), with additional remote contacts every 2 months and unscheduled visits as needed over a maximum of 30 months. IGRA testing was conducted at baseline and Month 12, with continuous surveillance for suspected TB. Baseline procedures included full physical exams, HIV, IGRA testing, and medical history review. Suspected TB cases (defined as participants presenting with unexplained cough, unexplained fever, night sweats, unintentional weight loss or hemoptysis) triggered unscheduled evaluations with up to three sputum sample collections, preferably within one week, for microbiological laboratory assessments and one blood sample collection for HIV, IGRA, and exploratory biomarkers.

### Laboratory Assessments

IGRA testing was conducted using the QFT-Plus assay (QuantiFERON®-TB Gold Plus, Qiagen, MD, USA) using modified manufacturer protocols for sample processing, specifically target goals for sample processing within four hours of sample collection and 20 hours incubation time for antigen stimulation [7]. This modified protocol was followed to reduce analytical variability [15-17]. QFT-Plus ELISA standards, controls, and analyses were conducted according to the manufacturer’s protocol.

For suspected TB visits, up to three sputum samples were collected preferably within one week, stored at 4°C, and shipped to the central laboratory within a target of seven days. At the central laboratory, samples were decontaminated (1% final concentration NaOH) and processed. Samples were tested using the Xpert® MTB/RIF Ultra assay (Xpert Ultra, Cepheid, Sunnyvale, CA) per manufacturer instructions and inoculated into the BACTEC MGIT culture system (Becton Dickinson, Franklin Lakes, NJ, USA). MGIT cultures with a positive signal were confirmed as *Mtb* using MPT64 antigen testing. Contaminated MGIT cultures were decontaminated and re-incubated. Final culture results were categorized as Negative, Contaminated (Inconclusive), Positive (MGIT+MPT64+), or Positive (MGIT+MPT64+) with Contamination.

### Exploratory Analysis Populations

For inclusion in this exploratory analysis, participants had to have QFT-Plus results obtained at the central laboratory, a valid IGRA result (i.e. not indeterminate), and transfer of all 4 quantitative values (NIL, TB1, TB2, Mitogen) from the QFT-Plus test at baseline (n=5,259 total). Participants from India were excluded from the exploratory analysis because quantitative QFT-Plus values were not available.

The control group is defined as participants who completed baseline and Month 12 visits and were not suspected of TB or diagnosed with TB during the study period.

The Suspected TB group (n=328) includes all participants who, during the study, developed signs or symptoms consistent with TB but for whom protocol-specified microbiological testing did not confirm disease. For this analysis, participants with more than two sequential suspected TB visits within a 90-day period were excluded to reduce potential site-level heterogeneity in clinical evaluation practices, particularly at sites with frequent repeat suspected TB assessments in the absence of corresponding increases in lab-confirmed TB. The Suspected TB group encompasses participants who triggered a suspected TB visit and had negative or contaminated laboratory results, as well as participants for whom study investigators initiated TB treatment despite the absence of microbiological confirmation (n=33, classified as clinical TB). Quantitative IFNγ concentrations did not differ between participants categorized as clinical TB and those in the broader suspected TB group (data not shown); therefore, they were analyzed together in subsequent analyses as they together represented study participants with symptoms but no microbiological confirmation.

The lab-confirmed TB group (n=23) is defined as participants with a microbiological confirmed TB diagnosis within the study period in the per-protocol population. Lab-confirmed TB is defined in the protocol as having at least one positive *Mtb* MGIT culture and/or at least one positive Xpert Ultra assay, based on 3 sputum samples collected at 3 different visits preferably within one week. Within the lab-confirmed TB group, we further differentiated based on the concordance of different tests and samples. The stringent TB case group (n=15) were defined as having two or more positive tests on one or more of the sputum samples collected during the suspected TB visit window. The single-positive TB case group (n=8) were defined as only having a single positive test (MGIT culture or Xpert Ultra) on only one of the sputum samples collected during the suspected TB visit window. For the Xpert Ultra assay, trace results were included in the positivity call in the TBV02-E01 primary analysis and thus were included in exploratory analyses as well (note: only three lab-confirmed TB cases in the exploratory population were determined using Xpert Ultra Trace results, two of which were classified as single-positive case and one that was used in combination with MGIT culture results to be classified as a stringent case [21]).

### Statistical Analysis

Comparisons of quantitative IFNγ concentrations across TB outcome groups were performed using the Kruskal-Wallis test to evaluate differences in medians. Subsequent pairwise tests between groups were done using the Mann-Whitney U test with Bonferroni correction applied within each set of tests (Supplementary Table 1). Spearman’s rank correlation coefficient (Spearman’s ρ) was used to assess associations between baseline quantitative IGRA values and the number of days to lab-confirmed TB diagnosis. All statistical analyses were conducted using the SciPy library in Python (version 1·15·2; https://scipy.org/).

TB1 and TB2 concentrations were first compared across TB outcome groups in the full exploratory population as all groups had participants who were IGRA positive or negative at baseline (Table 1). The QFT-Plus test qualitative output is determined by the manufacturer’s algorithm using NIL, TB1, TB2, and mitogen values, with positivity defined as a NIL-subtracted TB1 or TB2 value that is ≥0.35 International Units per milliliter (IU/ml) and >25% of the NIL value.^6^ By design, IGRA-negative individuals generally have low TB1 and TB2 IFNγ concentrations, potentially biasing analyses. To address this, we also completed an analysis restricted to IGRA-positive participants at baseline.

**Table 1:** Baseline Characteristics of the Exploratory Analysis Population. Demographics of comparator groups in the exploratory analysis population within the larger TB Epidemiology Study TBV02-E01. The lab-confirmed population is divided into two subsets, single positive case definition (single-positive) and stringent case definition (stringent). Means and standard deviations (SD) or percentage of exploratory analysis population are shown.

|  |  | Control | Suspected TB | Lab-Confirmed TB |  |
| --- | --- | --- | --- | --- | --- |
|  |  |  |  | Single-Positive Sputum Test | Stringent (≥2 Positive Sputum Tests) |
| N |  | 4908 | 328 | 8 | 15 |
| Age (Mean and SD) |  | 24.2 (4.9) | 24.7 (4.9) | 20.8 (4.7) | 27 (4.1) |
| Male |  | 46.3% | 52.4% | 25.0% | 53.3% |
| Race | Black | 55.3% | 77.1% | 87.5% | 46.7% |
|  | Asian | 37.0% | 15.2% | 12.5% | 46.7% |
|  | White | 2.2% | 3.7% | 0.0% | 0.0% |
|  | Multiple | 5.4% | 4.0% | 0.0% | 0.0% |
|  | Other | 0.2% | 0.0% | 0.0% | 6.7% |
| Healthcare Worker/Healthcare Student |  | 3.7% | 0.9% | 0.0% | 0.0% |
| Body Mass Index (BMI) (m/kg<sup>2</sup>) (Mean and SD) |  | 24 (5.6) | 23.4 (6.3) | 25.4 (9.0) | 21.3 (5.0) |
| Known BCG Vaccination |  | 79.2% | 50.3% | 50.0% | 66.7% |
| HIV Positive |  | 4.7% | 10.4% | 12.5% | 20.0% |
| IGRA+ at Baseline |  | 31.2% | 40.5% | 12.5% | 86.7% |
| Follow-up Duration, days (Mean [SD]) |  | 533 (86) | 592 (100) | 534 (179) | 525 (135) |
| Time to TB Diagnosis, days (Mean [SD]) |  | Not Applicable | Not Applicable | 320 (239) | 236 (133) |

### Evaluation of predictive value

The utility of quantitative IGRA values as predictive biomarkers for symptomatic TB disease development was evaluated using sensitivity, specificity and positive predictive values, which were calculated based on the ability to differentiate between pairs of TB outcome groups over the study follow-up period (listed in Table 1). Additionally, Youden’s J, F1-score and AUROC were calculated. ROC curves, positive predictive value and F1-score were calculated using scikit-learn in python (version 1·6·1; https://scikit-learn.org/).

## Results

This exploratory analysis included 5,259 participants from 13 countries: 4,908 in the control group, 328 in the suspected TB group, and 23 in the lab-confirmed TB group (15 meeting stringent case definitions and 8 meeting single-positive case definitions) (Table 1). Overall population characteristics were similar between the full study [21] and this exploratory analysis subset. Baseline characteristics were generally comparable across groups, although a higher proportion of people living with HIV and IGRA positivity were observed in the lab-confirmed TB group (stringent case definition) (Table 1). The mean follow-up duration in the study was comparable across groups (525-592 days, Table 1).

Using the full exploratory cohort, we analyzed the difference in baseline IFNγ concentrations between participants who did not progress to TB (controls) compared to participants who were later suspected of TB but with no laboratory confirmation and participants who later developed TB with lab confirmation. We intentionally included both IGRA-positive and IGRA-negative participants at baseline to allow us to evaluate QFT-Plus quantitative values as a continuous measure and assess their potential role as a one-time screening or stratification biomarker for clinical research applications in broad populations. At baseline, TB1 IFNγ concentrations were significantly higher in the suspected TB group compared to the control group (0.07 IU/mL vs. 0.03 IU/mL, p=0.0008) with a trend towards higher IFNγ levels in the lab-confirmed TB group compared to the control (1.38 IU/mL vs. 0.03 IU/mL, p=0.09) (Figure 1A). For TB2, IFNγ levels were significantly higher in both suspected and lab-confirmed TB groups relative to the control group (0.09 IU/mL vs. 0.04 IU/mL, p=0.01; 1.84 IU/mL vs 0.04 IU/mL, p=0.01, respectively) (Figure 1B).

**Figure 1.**
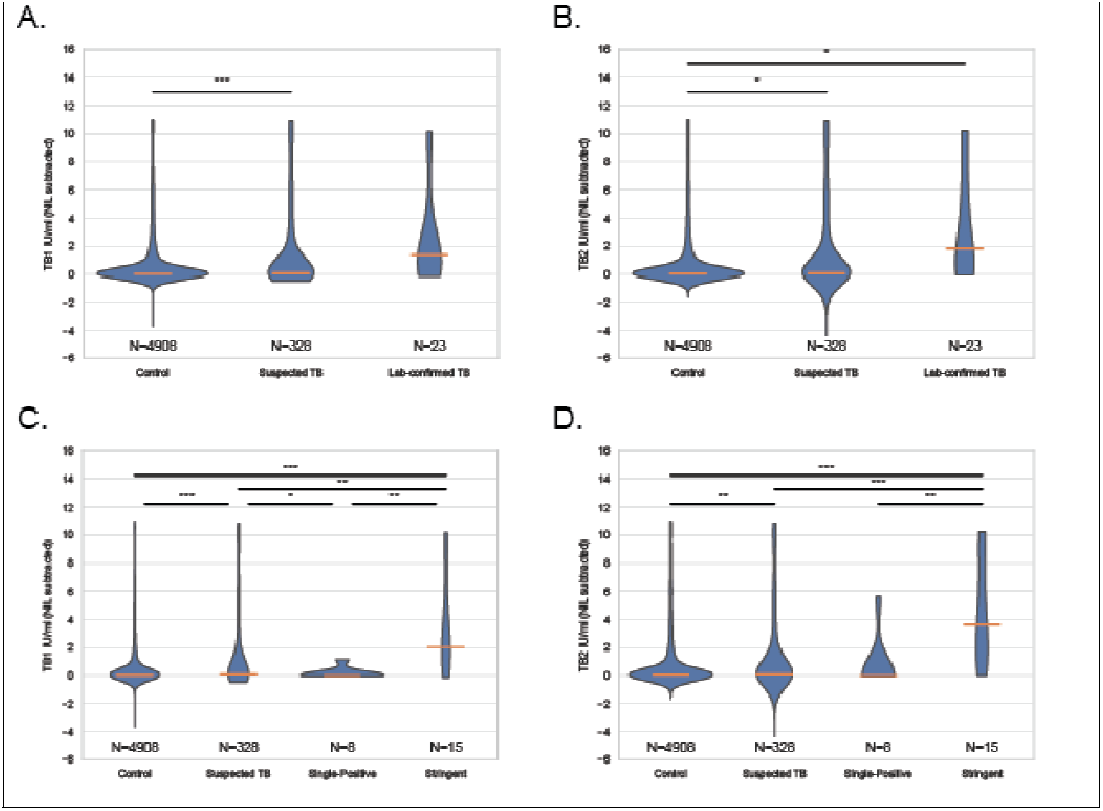
Baseline TB1 and TB2 IFN_γ_ Concentrations Among Participants by TB Outcome Classification in IGRA-Positive and –Negative Participants. (A–B) Violin plots depict IFN_γ_ concentrations (IU/ml, NIL-subtracted) from TB1 (A) and TB2 (B), antigen tubes measured at baseline among participants who did not develop TB (Control), were suspected of TB but not microbiologically confirmed (Suspected TB), or had lab-confirmed TB during follow-up. Statistical comparison by Kruskal-Wallis test showed significant differences across groups for all measures: TB1 (p=0.0001), TB2 (p=0.0001). (C-D) Box plots show baseline TB1 (C) and TB2 (D) IFN_γ_ levels stratified by more granular TB outcome groups: Control, Suspected TB, Single-assay Positive TB cases (1 positive culture or Xpert result), and Stringent TB Cases (≥2 positive microbiological results and/or sputum samples). Kruskal-Wallis tests were significant for TB1 (p=0.000002), TB2 (p=0.000001). Pairwise comparisons ^*^ = p< 0.05, ^**^ = p < 0.01, ^***^ = p < 0.001. Orange bar represents median value per group.

We next compared baseline IFNγ concentrations between groups in the full exploratory cohort using stringent and single-positive TB case definitions. The stringent case definition mirrors the primary endpoint being adopted in current and upcoming TB vaccine efficacy trials [22] and therefore provides a relevant benchmark for evaluating endpoint-matched biomarker performance. Compared to the control group, baseline TB1 and TB2 IFNγ concentrations were significantly higher in participants with suspected TB (TB1: 0.07 IU/mL vs. 0.03 IU/mL, p=0.001; TB2: 0.09 IU/mL vs. 0.04 IU/mL, p=0.02) and those meeting the stringent case definition (TB1: 2.02 IU/mL vs. 0.03 IU/mL, p=0.009; TB2: 3.59 IU/mL vs. 0.04 IU/mL, p=0.00003), but not in participants meeting single-positive case definition (Supplementary Table 1, Figure 1C, D). Participants meeting the stringent case definition had significantly higher TB1 and TB2 IFNγ concentrations than those meeting the single positive case definition (Supplementary Table 1, Figure 1C, D). Notably, median TB2 IFNγ concentration in participants meeting the stringent case definition (3.59 IU/mL, Figure 1D) was markedly higher when compared to the median of the broader lab-confirmed group (1.84 IU/mL, Figure 1B).

We then restricted the analysis to participants who were IGRA-positive at baseline to reduce bias from the low IFNγ values characteristic of IGRA-negative individuals and assess the biomarker’s potential utility in use cases limited to IGRA-positive populations. This reduced our TB cases as only 14/23 lab-confirmed participants were IGRA positive at baseline (n=13 stringent case definition, n=1 single-positive). In IGRA-positive participants, TB1 IFNγ concentrations were significantly higher in the suspected TB group compared to control (2.6 IU/mL vs. 1.64 IU/mL, p=0.004), but not in the lab-confirmed TB group (2.02 IU/mL vs. 1.64 IU/mL, p=0.3) (Figure 2A) or in participants meeting stringent (2.02 IU/mL vs. 1.64 IU/mL, p=0.5) or single-positive case definition (1.14 IU/mL vs. 1.64 IU/mL, p=0.4) (Figure 2C). For TB2, IFNγ concentrations were significantly higher in both the suspected TB and lab-confirmed TB groups relative to control group (2.79 IU/mL vs. 1.87 IU/mL, p=0.01; 4.74 IU/mL vs. 1.87 IU/mL, p=0.02) (Figure 2B), with higher levels in the smaller number of IGRA-positive participants meeting the stringent case definition although the difference was not significant (3.86 IU/mL vs. 1.87 IU/mL, p=0.09) (Figure 2D).

**Figure 2.**
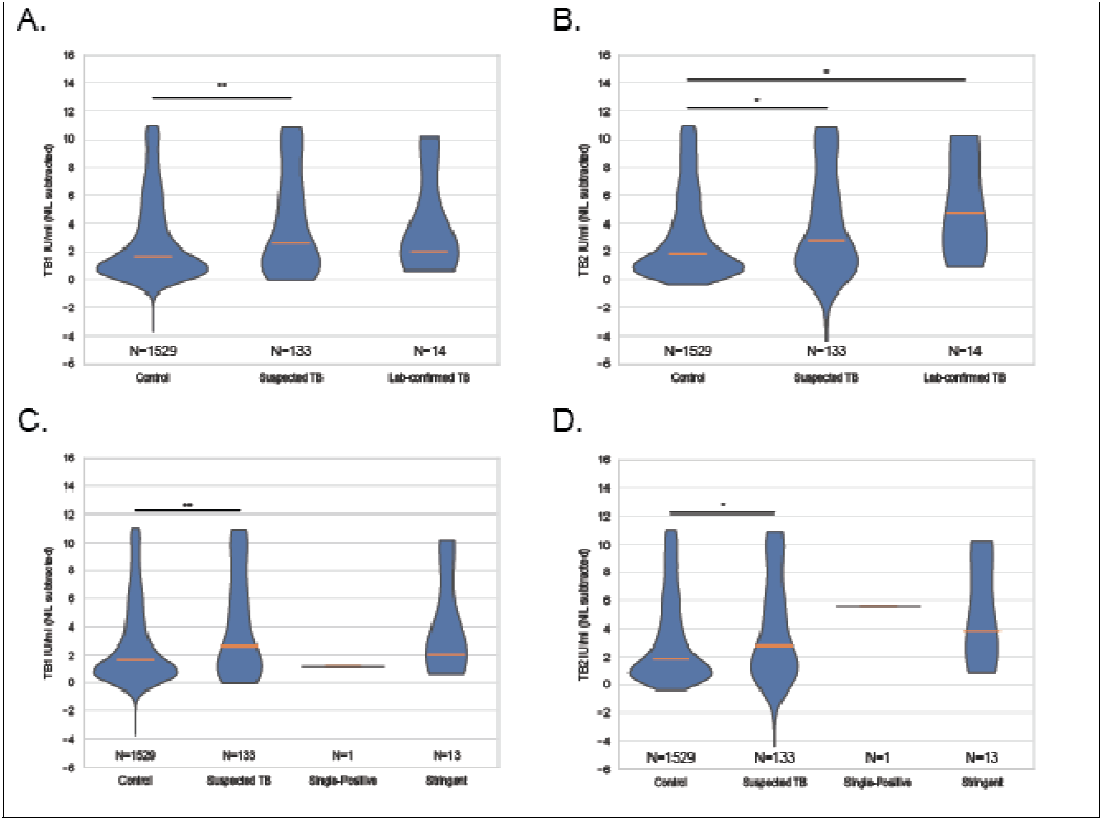
Baseline TB1 and TB2 IFN_γ_ Concentrations by TB Outcome Classification in IGRA-Positive Participants. (A–B) Violin plots show IFN_γ_ concentrations (IU/ml, NIL-subtracted) from TB1 (A) and TB2 (B) antigen tubes measured at baseline among IGRA-positive participants who did not develop TB (Control), were suspected of TB but not microbiologically confirmed (Suspected TB), or had lab-confirmed TB during follow-up. Kruskal-Wallis tests showed significant group differences for TB1 (p=0.002), TB2 (p=0.0008). (C-D) Box plots display TB1 (C) and TB2 (D) IFN_γ_ concentrations stratified by TB outcome groups: Control, Suspected TB, Single-assay positive TB cases (1 positive culture or Xpert result), and Stringent TB cases (≥2 positive microbiological results and/or sputum samples). Kruskal-Wallis tests showed significant group differences for TB1 (p=0.004), TB2 (p=0.002) Pairwise comparisons ^*^ = p< 0.05, ^**^ = p < 0.01, ^***^ = p < 0.001. Orange bar represents median value per group.

Analysis of correlation between IFNγ concentrations at baseline and time to diagnosis of lab-confirmed TB disease showed trends between baseline TB1 or TB2 IFNγ concentration and days to first positive laboratory test (TB1: r=-0.36, p=0.08; TB2: r=-0.34, p=0.10; Figure 3). Building on the observed associations between baseline IFNγ concentrations and subsequent TB outcomes, we next evaluated the sensitivity, specificity, and positive predictive value (PPV) of TB1 and TB2 measures. Given our interest in assessing the value of quantitative IFNγ measures as a single baseline biomarker for clinical research applications, including trial enrollment and stratification, we evaluated TB1 and TB2 IFNγ concentrations as a one-time measure independent of baseline IGRA status to predict TB development over the follow-up period. When differentiating laboratory-confirmed TB cases from controls, both markers showed similar performance in sensitivity (TB1: 52%; TB2: 57%) and specificity (TB1: 81%; TB2: 78%; (Table 2). Area under the Receiver Operating Characteristic curve (AUROC) values were also comparable across markers (TB1: 0.63; TB2: 0.67) (Figure 4A, B).

**Table 2:** Predictive Performance of Baseline IFN_γ_ Concentrations from QFT-Plus (TB1, TB2) for Progression to TB. Sensitivity, specificity, positive predictive value (PPV), and Youden’s J-derived optimal cutoff values are presented for each IFNγ measure across comparator groups. Analyses were conducted for two case definitions: (1) Lab-confirmed TB (n=23) versus controls (n=4,908) and suspected TB (n=328), and (2) Stringent case definition (n=15) versus controls and suspected TB. The Youden Cutoff represents the IFNγ threshold (in IU/mL) at which sensitivity and specificity are optimized.

| *Lab-confirmed* |  |  |  |  |  |  |
| --- | --- | --- | --- | --- | --- | --- |
|  | Group 1 | Group 2 | Youden Cutoff | Sensitivity | Specificity | PPV |
| TB1 | Control | Suspected TB | 0.50 | 36% | 75% | 8.6% |
|  | Control | Lab-Confirmed TB | 1.14 | 52% | 81% | 1.3% |
|  | Suspected TB | Lab-Confirmed TB | 0.65 | 57% | 68% | 11.1% |
| TB2 | Control | Suspected TB | 0.55 | 37% | 74% | 8.7% |
|  | Control | Lab-Confirmed TB | 0.89 | 57% | 78% | 1.2% |
|  | Suspected TB | Lab-Confirmed TB | 0.89 | 57% | 70% | 11.5% |
| *Stringent Case Definition* |  |  |  |  |  |  |
|  | Group 1 | Group 2 | Youden Cutoff | Sensitivity | Specificity | PPV |
| TB1 | Control | Suspected TB | 0.5 | 36% | 75% | 8.6% |
|  | Control | Stringent Case | 0.65 | 80% | 77% | 1.0% |
|  | Suspected TB | Stringent Case | 0.65 | 80% | 68% | 10.3% |
| TB2 | Control | Suspected TB | 0.55 | 37% | 74% | 8.7% |
|  | Control | Stringent Case | 0.89 | 80% | 78% | 1.1% |
|  | Suspected TB | Stringent Case | 0.89 | 80% | 70% | 10.7% |

**Figure 3:**
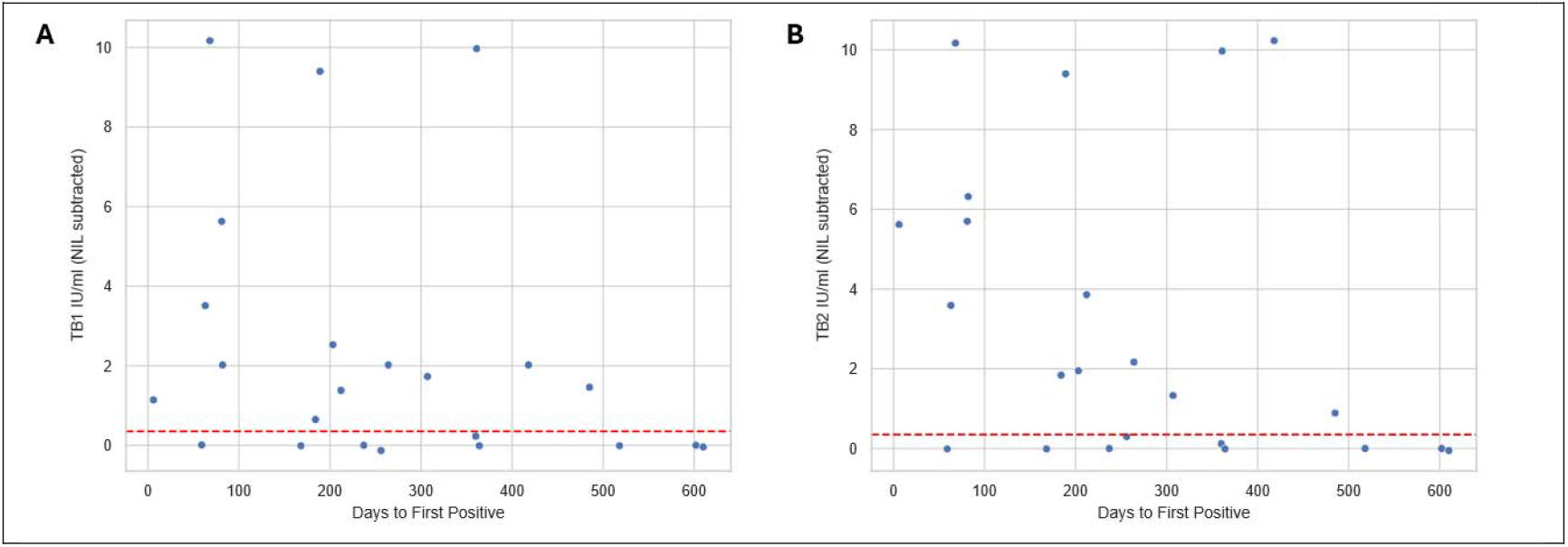
Correlation between Baseline IFN_γ_ Concentrations and Time to TB Diagnosis among Lab-Confirmed TB Cases. Scatter plots show baseline IFN-_γ_ concentrations (NIL-subtracted, in IU/mL) for (A) TB1 and (B) TB2 plotted against the number of days from baseline to first positive TB laboratory test. Spearman correlation coefficients (ρ) and p-values for (A) ρ=-0.36, p=0.08 and (B) ρ=-0.34, p=0.10. Dashed red lines indicate the manufacturer’s recommended positivity threshold of 0.35 IU/mL

**Figure 4:**
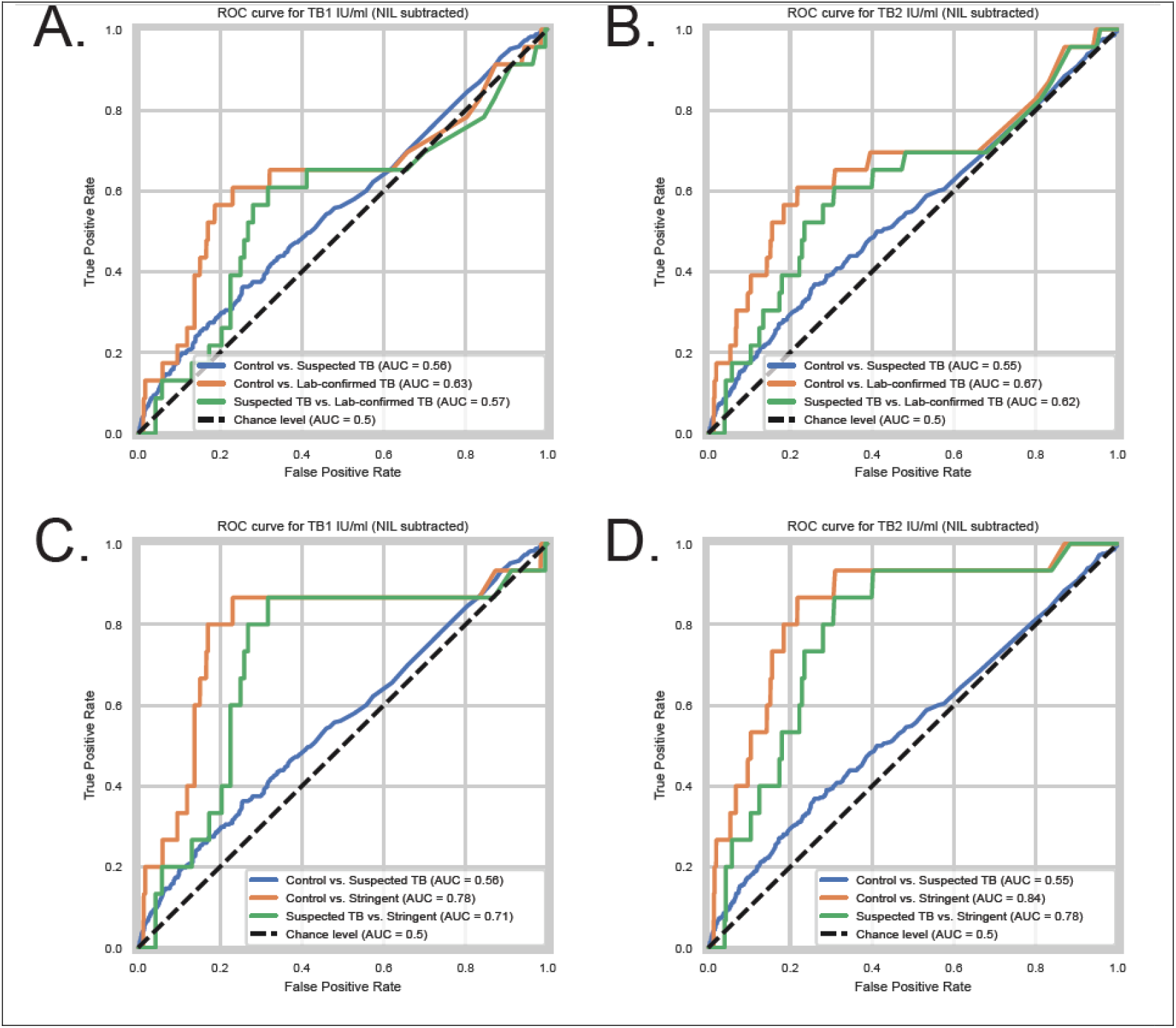
Receiver Operating Characteristic (ROC) Curve Analyses of Baseline IFN_γ_ Concentrations for Predicting TB Disease Progression. Panels A–B depict ROC curves for distinguishing outcome groups using baseline IFN_γ_ levels from (A) TB1 and (B) TB2 in the analysis based on lab-confirmed TB cases (n=23). Comparisons include Control vs. Suspected TB (blue), Control vs. Lab-Confirmed TB (orange), and Suspected TB vs. Lab-Confirmed TB (green). Panels C-D show ROC curves for the same IFN_γ_ measures, (D) TB1 and (E) TB2 in the stringent case definition subgroup (n=15). Comparisons include Control vs. Suspected TB (blue), Control vs. Stringent (orange), and Suspected TB vs. Stringent (green). Area under the curve (AUC) values are indicated for each comparison. Diagonal dashed lines indicate the line of no discrimination (AUROC = 0.5).

Performance improved further when applying the more stringent case definition and distinguishing against controls, with similar sensitivity (TB1: 80%; TB2: 80%) and specificity (TB1: 77%, TB2: 78%), exceeding the minimal WHO-recommended test thresholds for a predictive test for progression to TB disease.^19^ AUROC values were highest for TB2 (0.84) (Figure 4D). The PPV for progression to TB disease within two years was low across all measures (Table 2). Youden’s J values for distinguishing stringent TB cases from controls were higher than the standard QFT-Plus diagnostic cutoff of ≥0.35 IU/mL, with values of 0.65 IU/mL (TB1) and 0.89 IU/mL (TB2), suggesting that more strict cutoffs outside the diagnostic uncertainty zone [16] may also improve predictive accuracy to symptomatic disease.

## Discussion

We observed that independent of IGRA status, high baseline TB2 IFNγ concentrations from the QFT-Plus test were associated with subsequent progression to laboratory-confirmed symptomatic TB disease and were most elevated in IGRA-positive and -–negative participants meeting a stringent TB case definition. Association of baseline TB1 IFNγ concentration and eventual symptomatic disease progression was found only when including both including IGRA-positive and –negative individuals and in participants classified under a stringent TB case definition. However, consistent with previous studies [12-14], we found substantial variability and considerable overlap in IFNγ concentrations between participants who did and did not progress to symptomatic TB disease as well as no significant association between baseline measurements and time to laboratory confirmed TB. In addition, the relatively limited number of TB cases negatively impacted the observed PPV of these measures. Although promising, these observations highlight the current limitations of using QFT-Plus quantitative IFNγ measurements alone as reliable predictive biomarkers to symptomatic TB disease.

When considering the dynamic TB spectrum, increasing TB-specific IFNγ concentrations may reflect evolving host-*Mtb* interactions, with rising mycobacterial burden resulting in increased TB-specific T-cell activation and effector responses [8, 23-25]. The elevated IFNγ responses observed in this state could signal early breakdown of immune containment, preceding progression to symptomatic disease. As no direct measurement of the presence of TB was performed at baseline in this study, we were unable to unequivocally identify or exclude asymptomatic TB among participants in this analysis. Future studies specifically incorporating populations with asymptomatic TB are needed to determine how TB1 and TB2 IFNγ concentrations perform across the full TB disease spectrum. For clinical or research applications in which distinguishing asymptomatic TB from other disease states is not required, QFT-Plus quantitative measures may still provide added value.

In this study, baseline TB2 IFNγ concentrations were progressively higher among participants in the suspected TB group (without microbiological confirmation), the lab-confirmed TB group (at least 1 positive microbiological test), and those meeting a stringent case definition (at least two positive microbiological tests). This gradient suggests that the strength of association between TB2 IFNγ concentrations and TB outcomes increases with greater diagnostic certainty. Only when including participants meeting the stringent case definition did both TB1 and TB2 IFNγ measures exceed the WHO minimum test characteristics (>75% sensitivity, >75% specificity) for predicting progression to symptomatic TB [4]. These findings have practical implications for clinical research settings where a biomarker may be needed as a single baseline screening tool to enrich or stratify participants in vaccine efficacy trials or prevention studies. Notably, recent TB vaccine trials have adopted stringent case definitions as their primary efficacy endpoint. Under these conditions, quantitative IFNγ concentrations, particularly TB2 values, may offer meaningful predictive or stratification utility beyond the binary classification afforded by qualitative QFT-Plus result. These findings also highlight that with continuous efforts to refine endpoint definitions for TB vaccine and therapeutics trials [26], candidate biomarkers should be re-evaluated to ensure that their performance aligns with the specificity of the selected endpoints, thereby minimizing the risk of dismissing potentially valuable markers that underperform under broader case definitions.

This study has several limitations. As an exploratory analysis of data from a large epidemiological study, it was not specifically powered to assess the predictive performance of quantitative IFNγ concentrations from the QFT-Plus assay. Furthermore, the limited number of TB cases precluded robust subgroup analyses that would help determine the generalizability and stability of these associations across subsets. However, the data generated provides an important foundation for building power calculations to inform the design of future studies which can rigorously assess the utility of these measures. Finally, because the study focused on high-burden TB settings to support phase 3 vaccine trial readiness, the findings may not apply to low incidence settings or general populations.

In conclusion, QFT-Plus IFNγ concentrations, and TB2 values in particular, may be informative measures as early indicators of symptomatic TB disease progression. While limited in standalone predictive value, the associations between these biomarkers in individuals who progress to symptomatic TB disease as defined under the stringent TB case definition are promising for future clinical research applications, especially considering the frequent use of the QFT-Plus assay in TB vaccine trials, which makes these data readily accessible. Integrating QFT-Plus quantitative IFNγ readouts with additional measures may yield more robust composite biomarkers for identifying individuals with the highest risk. For clinical research, such biomarkers could enable enrichment of trial populations with higher event rates, improve power for detecting efficacy, and align risk prediction more closely with the symptomatic TB endpoints used in modern phase 3 TB vaccine trials.

### Gates MRI TB Epi Study Group

Q Bhorat (Soweto Clinical Trials Centre, Johannesburg), M Loveday (Botha’s Hill CRS, Durban, South Africa), C Duran Palma (Trident Clinical, Homestead, South Africa), J Lombaard (Josha Research, Mangaung, South Africa), S Cossie (CRISMO Bertha Gxowa Research Centre, Ekurhuleni, Gauteng, South Africa), JG Geldenhuys (FCRN Clinical Trial Centre Pty Ltd, Three Rivers Vereeniging, Gauteng, South Africa) K Ahmed (Setshaba Research Centre, Soshanguve, Gauteng, South Africa) E Spooner (PHOENIX Pharma (Pty) Ltd, HIV Prevention Research Unit, Ethekwini, Kwazulu – Natal, South Africa), L Tina (Victoria Biomedical Research Institute, US Army Medical Research Unit, Kombewa Clinical Research Centre, Kisumu, Western, South Africa), B Ogutu (Strathmore University, Centre for Clinical Research, Kisumu, Western, South Africa),R McClelland (Ganjoni Municipal Communicable Diseases Control Centre, Mombasa, Kenya), W Kilembe (Center for Family Health Research in Zambia (CFHRZ) – Lusaka, Zambia Emory HIV Research Project Lusaka, Lusaka, Zambia), S Kwame (Zambart University of Zambia, Zambart, RidgewayCampus, Lusaka, Zambia), M Inambao (Center for Family Health Research in Zambia (CFHRZ) – Ndola, Northrise, Zambia), E Sprinz (Hospital de Clinicas de Porto Alegre (HCPA) – PPDS, Floresta Porto Alegre – RS, Porto Alegre, Rio Grande do Sul, Brazil), E Cadena (Philippine Tuberculosis Society Inc, Quezon City, National Capital Region, Philippines), JR Gonong (Lung Center of The Philippines, Quezon City, Philippines), EJ Berame (Healthlink Iloilo, Iloilo City, Iloilo, Philippines), MGD Isidro (West Visayas State University Medical Center, Iloilo City, Iloilo, Philippines), G Zabat (Health Cube Medical Clinics, Quezon City, Philippines), RGM Veto (Tropical Disease Foundation, Makati City, Philippines), Javier R Lama (Asociación Civil Impacta Salud y Educación, Lima, Peru), E Sanchez (Hospital Nacional Sergio E. Bernales, Lima, Peru), V Nankabirwa (Makerere University - School of Public Health, Kampala, Uganda), HL Nguyen (Pham Ngoc Thach Hospital, Phường Số 12 Quận 5, Vietnam), VN Nguyen (National Lung Hospital, Hanoi, Vietnam), G Marks (Woolcock Institute Of Medical Research, Hanoi, Vietnam), B Alisjahbana (Universitas Padjadjaran, Bandung, Jawa Barat, Indonesia), Burhan E (Persahabatan Hospital, Jakarta, Indonesia), Kaswandani N (Puskesmas Kecamatan Kramat Jati, Jakarta, Indonesia), G Luyeye Matondo Mandiangu (Centre de Sante Maternite Binza, Ngaliema, Kinshasa, Democratic Republic of the Congo), HZ Swah Koko (Centre de Sante 2eme Rue, Limete, Kinshasa, Democratic Republic of the Congo), E Mwamba Kabunda (Hopital General de reference de Kabinda, Lingwala, Kinshasa, Democratic Republic of the Congo) S Viegas (Instituto Nacional de Saude, Marracuene, Mozambique), AL Garcia-Basteiro (Centro de Investigação em Saúde de Manhiça (CISM), Maputo, Mozambique)

## Acknowledgements

We are grateful to the study participants and their families for taking part in this trial and all clinical investigators, independent ethics committee members, site staff, PPD staff, and BARC staff for their contributions. We would like to thank all members of the TBV02-E01 Study Group including clinical operations. This study was funded by the Gates Foundation [INV-008522]. The conclusions and opinions expressed in this work are those of the authors alone and shall not be attributed to the Foundation. Under the grant conditions of the Foundation, a Creative Commons Attribution 4.0 License has already been assigned to the Author Accepted Manuscript version that might arise from this submission. Medical writing assistance was provided by Madeeha Aqil, PhD, CMPP™ and was funded by the Gates Medical Research Institute.

## Contributions

JS and MS were involved in strategy and data analysis. AFD and ACS were involved in the conception or design of the trial. JS, MS, AFD, LLH, AC, DG, AAH, MTG, SRH, WAH, JRL, MM, SM, VN, VCR, TR, JS, CK, AW, TMW, contributed to data collection or data generation. JS, MS, AFD, ACS, AC, and NF contributed to data interpretation. MS accessed and verified the data. JS and NF were responsible for the decision to submit the manuscript. All authors reviewed the draft critically, approved the final version to be submitted, and took accountability for all aspects of the published work.

## Data sharing

Anonymized participant-level data may be shared with external researchers in accordance with the study participants’ written and executed informed consent document and any local or applicable regulations on data sharing. Qualified researchers may submit a request for anonymized participant-level data along with a research proposal to Gates MRI’s Data Protection Officer at for review. The types of supporting information that could be shared with external researchers include: the Study Protocol, Statistical Analysis Plan, Informed Consent Form, Clinical Study Report, and analytic code. A data sharing agreement must be in place before any study data are shared. There are additional circumstances that may prevent the sharing of data with external researchers, including but not limited to contractual obligations to existing partners and any restrictions imposed by regulatory bodies.

