## Supplement table for "Associations Between QuantiFERON-TB Gold Plus IFN_γ_ Concentrations and Progression to Symptomatic Tuberculosis in Global High-Burden TB Settings"

| Group 1 | Group 1 N | Group 1 median | Group 1 IQR | Group 2 | Group 2 N | Group 2 median | Group 2 IQR | statistic | Median difference | p-value | q-value |
|---|---|---|---|---|---|---|---|---|---|---|---|
| *TB1 IFN gamma concentrations differences between controls, suspected TB, and Lab confirmed TB groups* | | | | | | | | | | | |
| Suspected TB | 328 | 0.07 | 1.73 | Control | 4908 | 0.03 | 0.53 | 901127.5 | -0.04 | 0.000277 | 0.00083 |
| Lab-confirmed TB | 23 | 1.38 | 2.275 | Control | 4908 | 0.03 | 0.53 | 71156 | -1.35 | 0.03048 | 0.091439 |
| Lab-confirmed TB | 23 | 1.38 | 2.275 | Suspected TB | 328 | 0.07 | 1.73 | 4285.5 | -1.31 | 0.274722 | 0.824165 |
| *TB2 IFN gamma concentrations between controls, suspected TB, and Lab confirmed TB groups* | | | | | | | | | | | |
| Suspected TB | 328 | 0.09 | 1.6825 | Control | 4908 | 0.04 | 0.62 | 882270.5 | -0.05 | 0.003467 | 0.0104 |
| Lab-confirmed TB | 23 | 1.84 | 5.66 | Control | 4908 | 0.04 | 0.62 | 75347.5 | -1.8 | 0.00544 | 0.016319 |
| Lab-confirmed TB | 23 | 1.84 | 5.66 | Suspected TB | 328 | 0.09 | 1.6825 | 4662 | -1.75 | 0.058321 | 0.174964 |
| *TB1 IFN gamma concentrations differences between controls, suspected TB, single-positive and stringent groups* | | | | | | | | | | | |
| Stringent | 15 | 2.02 | 3.15 | Control | 4908 | 0.03 | 0.53 | 57493.5 | -1.99 | 0.000164 | 0.000981 |
| Suspected TB | 328 | 0.07 | 1.73 | Control | 4908 | 0.03 | 0.53 | 901127.5 | -0.04 | 0.000277 | 0.00166 |
| Stringent | 15 | 2.02 | 3.15 | Single-Positive | 8 | 0 | 0.075 | 104.5 | -2.02 | 0.004449 | 0.026697 |
| Stringent | 15 | 2.02 | 3.15 | Suspected TB | 328 | 0.07 | 1.73 | 3507 | -1.95 | 0.005259 | 0.031553 |
| Suspected TB | 328 | 0.07 | 1.73 | Single-Positive | 8 | 0 | 0.075 | 1845.5 | -0.07 | 0.049185 | 0.29511 |
| Single-Positive | 8 | 0 | 0.075 | Control | 4908 | 0.03 | 0.53 | 13662.5 | 0.03 | 0.136041 | 0.816243 |
| *TB2 IFN gamma concentrations differences between controls, suspected TB, single-positive and stringent groups* | | | | | | | | | | | |
| Stringent | 15 | 3.59 | 6.275 | Control | 4908 | 0.04 | 0.62 | 61869 | -3.55 | 0.000005 | 0.00003 |
| Stringent | 15 | 3.59 | 6.275 | Suspected TB | 328 | 0.09 | 1.6825 | 3837.5 | -3.5 | 0.000242 | 0.001451 |
| Suspected TB | 328 | 0.09 | 1.6825 | Control | 4908 | 0.04 | 0.62 | 882270.5 | -0.05 | 0.003467 | 0.020801 |
| Stringent | 15 | 3.59 | 6.275 | Single-Positive | 8 | 0 | 0.04 | 105.5 | -3.59 | 0.003634 | 0.021802 |
| Suspected TB | 328 | 0.09 | 1.6825 | Single-Positive | 8 | 0 | 0.04 | 1799.5 | -0.09 | 0.072404 | 0.434422 |
| Single-Positive | 8 | 0 | 0.04 | Control | 4908 | 0.04 | 0.62 | 13478.5 | 0.04 | 0.124438 | 0.746628 |
| *TB1 IFN gamma concentrationsdifferences between controls, suspected TB, and Lab confirmed TB groups among baseline IGRA+ subjects* | | | | | | | | | | | |
| Suspected TB | 133 | 2.6 | 5.5 | Control | 1529 | 1.64 | 3.82 | 118425.5 | -0.96 | 0.001607 | 0.004822 |
| Lab-confirmed TB | 14 | 2.02 | 3.5725 | Control | 1529 | 1.64 | 3.82 | 13376 | -0.38 | 0.107323 | 0.321969 |
| Lab-confirmed TB | 14 | 2.02 | 3.5725 | Suspected TB | 133 | 2.6 | 5.5 | 975 | 0.58 | 0.774065 | 2.322195 |
| *TB2 IFN gamma concentrations differences between controls, suspected TB, and Lab confirmed TB groups among baseline IGRA+ subjects* | | | | | | | | | | | |
| Suspected TB | 133 | 2.79 | 5.37 | Control | 1529 | 1.87 | 4.17 | 116622.5 | -0.92 | 0.004879 | 0.014636 |
| Lab-confirmed TB | 14 | 4.74 | 6.625 | Control | 1529 | 1.87 | 4.17 | 15006.5 | -2.87 | 0.009519 | 0.028558 |
| Lab-confirmed TB | 14 | 4.74 | 6.625 | Suspected TB | 133 | 2.79 | 5.37 | 1149.5 | -1.95 | 0.150273 | 0.45082 |
| *TB1 IFN gamma concentrations differences between controls, suspected TB, single-positive and stringent groups among baseline IGRA+ subjects* | | | | | | | | | | | |
| Suspected TB | 133 | 2.6 | 5.5 | Control | 1529 | 1.64 | 3.82 | 118425.5 | -0.96 | 0.001607 | 0.009644 |
| Stringent | 13 | 2.02 | 3.9 | Control | 1529 | 1.64 | 3.82 | 12764.5 | -0.38 | 0.077165 | 0.462991 |
| Stringent | 13 | 2.02 | 3.9 | Single-Positive | 1 | 1.14 | 0 | 12 | -0.88 | 0.212825 | 1.276948 |
| Suspected TB | 133 | 2.6 | 5.5 | Single-Positive | 1 | 1.14 | 0 | 92 | -1.46 | 0.518063 | 3.108379 |
| Stringent | 13 | 2.02 | 3.9 | Suspected TB | 133 | 2.6 | 5.5 | 934 | 0.58 | 0.635405 | 3.812432 |
| Single-Positive | 1 | 1.14 | 0 | Control | 1529 | 1.64 | 3.82 | 611.5 | 0.5 | 0.729884 | 4.379306 |
| *TB2 IFN gamma concentrations differences between controls, suspected TB, single-positive and stringent groups among baseline IGRA+ subjects* | | | | | | | | | | | |
| Suspected TB | 133 | 2.79 | 5.37 | Control | 1529 | 1.87 | 4.17 | 116622.5 | -0.92 | 0.004879 | 0.029272 |
| Stringent | 13 | 3.86 | 7.45 | Control | 1529 | 1.87 | 4.17 | 13815.5 | -1.99 | 0.015317 | 0.091904 |
| Stringent | 13 | 3.86 | 7.45 | Suspected TB | 133 | 2.79 | 5.37 | 1060.5 | -1.07 | 0.179163 | 1.074977 |
| Single-Positive | 1 | 5.62 | 0 | Control | 1529 | 1.87 | 4.17 | 1191 | -3.75 | 0.334786 | 2.008718 |
| Suspected TB | 133 | 2.79 | 5.37 | Single-Positive | 1 | 5.62 | 0 | 44 | 2.83 | 0.569523 | 3.417136 |
| Stringent | 13 | 3.86 | 7.45 | Single-Positive | 1 | 5.62 | 0 | 6 | 1.76 | 1 | 6 |
